# Exacerbation of mitochondrial fission in human CD34^+^ cells halts erythropoiesis and hemoglobin biosynthesis

**DOI:** 10.1101/2020.07.31.230961

**Authors:** Alvaro M. Gonzalez-Ibanez, Lina M. Ruiz, Erik Jensen, Cesar A. Echeverria, Valentina Romero, Linsey Stiles, Orian Shirihai, Alvaro A. Elorza

## Abstract

Erythropoiesis is the most powerful cellular differentiation and proliferation system, with a production of 10^11^ cells per day. In this fine-tuned process, the hematopoietic stem cells (HSCs) generate erythroid progenitors, which proliferate and mature into erythrocytes. During erythropoiesis, mitochondria are reprogrammed to drive the differentiation process before finally being eliminated by mitophagy. In erythropoiesis, mitochondrial dynamics (MtDy) is expected to be a regulatory key point that has not been described previously. We described that a specific MtDy pattern is occurring in human erythropoiesis from EPO-induced human CD34^+^ cells, characterized by a predominant mitochondrial fusion at early stages followed by predominant fission at late stages. The fusion protein MFN1 and the fission protein FIS1 are shown to play a key role in the accurate progression of erythropoiesis. Fragmentation of the mitochondrial web by the overexpression of FIS1 (gain of fission) resulted in both the inhibition of hemoglobin biosynthesis and the arrest of erythroid differentiation, keeping cells in immature differentiation stages. These cells showed specific mitochondrial features as compared with control cells, such as an increase in round and large mitochondria morphology, low mitochondrial membrane potential and a drop in the expression of the respiratory complexes II and IV. Interestingly, treatment with the mitochondrial permeability transition pore (mPTP) inhibitor cyclosporin A, rescued mitochondrial morphology, hemoglobin biosynthesis and erythropoiesis. Studies presented in this work revealed MtDy as a hot spot in the regulation of erythroid differentiation which might be signaling downstream for metabolic reprogramming through the aperture/close of the mPTP.

**Key Points:**

-. Excessive fission disrupts erythroid progression, heme biosynthesis and mitochondrial function, keeping cells mostly in progenitors and proerythroblast stage.
-. Mitochondrial Dynamics signaling for erythroid differentiation involves FIS1 and the mPTP

## 1. Introduction

Mitochondria play several critical roles throughout erythroid differentiation to produce red blood cells. In early expansion of erythropoiesis -from HSC to CFU-E-, mitochondria are involved in stemness and stem cell differentiation, regulating energy metabolism and metabolic reprogramming from glycolytic and proliferative, to oxidative and differentiating metabolism^1–6^. In late expansion -from proerythroblasts to polychromatophilic erythroblasts-, mitochondria are involved in iron metabolism and heme biosynthesis. ^1,2^. Terminal erythroid maturation -from orthochromatic erythroblasts to mature erythrocytes- is mainly distinguished by mitochondria driving their own elimination through mitophagy to produce a fully mature erythrocyte ^3,4^.

Mitochondrial dynamics (MtDy) refers to continuous fission and fusion events which participate in mitochondrial turnover ˝coupled to mitophagy- and cell signaling. These interactions, in terms of frequency and remodeling of the mitochondrial web, reflect mitochondrial adaptive responses to accomplish cell proliferation, differentiation, energy demand, stress response, cell survival and death ^3–5^. Mitochondrial fission is dependent on the protein DRP1 that has been recognized as a mitochondrial fission promoter ^6,7^. It is located in the cytosol, but translocases to mitochondria to bind the mitochondrial receptor proteins FIS1, MFF, MiD49 and MiD51 ^8,9^, which are all located in the mitochondrial outer membrane (MOM). Mitochondrial fusion is dependent on MFN1 and MFN2, also located in the MOM; and OPA1, which is located in the mitochondrial inner membrane ^10,11^.

Mitochondria in stem cells feature low biomass and mtDNA copy number presenting immature cristae ^12^. The OXPHOS capacity and ROS generation are minimal, and the mitochondrial membrane potential is mainly sustained by the ATP synthase in reverse mode i.e. consuming glycolysis-made ATP for protons translocation. Upon differentiation, mitochondria start fusing and increasing their respiratory capacity, mitochondrial membrane potential and ROS generation; switching from a glycolytic to oxidative metabolism. ^12–17^. In this regard, MtDy act as modulators of stemness and stem cell fate decisions. It has been shown that deregulation of the mitochondrial fission protein DRP1 was associated with a loss of pluripotent markers (Nanog, Oct4 and Ssea) in mouse embryonic stem cells ^18^. Furthermore, cell differentiation from murine mesenchymal stem cells into osteo, chondro, and adipocytes requires specific MtDy changes. In adipo and osteogenesis, mitochondrial elongation and expression of MFN1 and MFN2 are increased in contrast to chondrogenesis, where mitochondrial fragmented morphology and expression of DRP1 and FIS1 are favored ^19^. The right mitochondrial fusion/fission ratio has been also described in the immune system by controlling macrophage migration ^20^; the antiviral signaling^21^ and the T-cell fate through metabolic reprogramming ^22^.

Erythropoiesis is the only system in nature capable of producing more than 100 billion cells per day starting from stem cells. This system involves exceptional high rates of proliferation and differentiation where the role of mitochondrial dynamics has not been studied. Our group has previously shown that exposure to non-cytotoxic copper overload or deficiency in erythropoietic cells modifies cell proliferation and differentiation along with changes in mitochondrial morphology, expression of MtDy proteins and the rate of mitochondrial fission and fusion events ^23,24^. In this work, we hypothesized that a specific pattern of MtDy is needed for the appropriate commitment and differentiation of HSC into the red cell lineage.

## 2. Methods

### Molecular tools

pLVCTH-shFIS1 and pLVCTH-shMFN1were used for gene silencing and pWPI FIS1 and pWPI MFN1, for gene overexpression. Lentiviral particles were produced in HEK-293T cells with the vectors psPAX2 (12259, Addgene), pMD2.G (12259, Addgene) and either pLVCTH or pWPI.

### Cell isolation, culture and lentiviral transduction

Mouse G1E-ER cells were cultured and differentiated to red cells according to ^25^ Primary human CD34+ cells were obtained from umbilical cord blood, after a normal full-term delivery (informed consent was given) as described in ^24^ and culture according to ^26^. For transduction, 5×10^4^ cells were seeded a 96-well plate with 8 μg/mL Polybrene (107689, Sigma) and exposed overnight to lentiviral particles.

### Erythroid Progression

Erythroid progression was followed by immunostaining with phycoerythrin (PE)–conjugated anti-human CD235a (1:100) (116207, BioLegend) and Allophycocyanin (APC)–conjugated anti-CD71 (1:100) (551374, BD Pharmingen). Flow cytometry was carried out on a BD Accuri C6 Cytometer (BD Biosciences). In addition, 3,3,5,5-tetramethylbenzidine (Sigma-Aldrich) was used for benzidine staining according to ^27^

### Real-time qPCR

Total RNA was isolated with the PureLink RNA Mini Kit (12183018A, Thermo) and cDNA was synthesized with ProtoScript First Strand cDNA (E6300S, New England Biolabs). RT-PCR was performed with the Power SYBR Green PCR Master Mix 2X (4367659, Thermo). Primers are listed in the Supplementary Table I.

### Immunoblotting

Cells were lysed in RIPA buffer with protease inhibitor complex HALT 1X (78420, Roche). Total proteins were run on SDS-PAGE, transferred to PVDF membranes and blotted with the antibodies against FIS1 (ab71498, Abcam), MFN1 (ab104274,Abcam), ß-Actin (ab8227, Abcam), Caspase-3 (9665, Cell Signaling), Caspase-8 (sc-5263, Santa Cruz), Caspase-9 (sc-8355, Santa Cruz), HSP70 (TA309356, Origene) and Total OXPHOS Human WB Antibody Cocktail (ab110411, Abcam).

### Immunofluorescence

Cells were attached on Poly-L-lysine (P4707, Sigma) coated slides, fixed with 4% paraformaldehyde (15711, Electron Microscopy Sciences) and permeabilized with 0,1% Triton X-100. All cells washes were in PBS/1% BSA. The primary Anti-VDAC1 (ab15895, Abcam) and the secondary Alexa Flour 594 (A11037, Life) antibodies were used. Cells were mounted with DAPI Fluoromount-G (0100-20, Southern Biotech) and examined with a Leica TCS LSI confocal microscope.

### Mitochondria staining

For mitochondrial morphology, cells were stained with 10nM MitoTracker Red CMXRos (M7512, Life), fixed in 4% paraformaldehyde (15711, Electron Microscopy Sciences) and mounted with DAPI Fluoromount-G (0100-20, Southern Biotech). For membrane potential, cells were stained with 10 nM TMRE (ab113852,Abcam) and visualized by FACS and live confocal imaging.

### Transmission Electron Microscopy

Samples were processed according to the TEM facility from P. Catholic University of Chile. Ultrathin sections of 60-70 nm were viewed with Philips Tecnai 12 a 80 kV.

### Mitochondrial morphometric analysis

Z-slides from TEM and confocal microscopy were used for mitochondrial circularity, perimeter and area measurements obtained with the FIJI-imageJ software ^28^. Scatter plots with histograms were done with the Python-based software Matplotlib 3.3.0.

### Statistical analysis

Statistical analysis was done with the GraphPad Prism 7 (GraphPad Software Inc) with a significance level α < 0.05. ANOVA and Bonferroni post hoc test were performed to compare averages between groups. Linear regression analysis was performed to compare slopes. R^2^ and the Mean Square Error (MSE, the lower the MSE the higher the accuracy of prediction) were also calculated.

## 3 Results

### 3.1 Mitochondrial Dynamics and Morphology in Erythropoiesis

Mitochondrial dynamics and morphology were first investigated in the mouse erythropoietic cell line G1E-ER (G1E cells expressing a GATA-1 construct fused to an estrogen receptor ligand-binding domain), given their fast and synchronic erythroid differentiation when stimulated with ß-estradiol. Mitochondrial morphology and the expression of MtDy genes during erythroid differentiation was examined by confocal microscopy and quantitative PCR respectively. 24 hrs. post ß-estradiol-induced erythropoiesis, mitochondrial web changed from elongated to fragmented (Fig. S1A). The mRNA expression pattern of *fis1*, *opa1*, *mfn1*, and *mfn2* genes was measured over 48 hrs. post ß-estradiol-induced erythropoiesis at regular 12 hrs. intervals by quantitative real time PCR. The mRNA concentration of *fis1*, which ranged from 2.27 fM to 5.82 fM, was considerably greater than those of the *opa1*, *mfn1*, and *mfn2* fusion genes which were all under 1 fM at every time point. It was also observed that *mfn1* transcripts were tenfold more abundant than *mfn2* ones (Fig. S1B). While the gene expression of *opa1* and *mfn2* seems not to change over mouse erythroid differentiation, *fis1* and *mfn1* genes showed a specific and differential mRNA expression pattern, progressively increasing their expression until 36 hrs. to decline later at 48 hrs. post ß-estradiol treatment. (Fig. S1B). These results suggest *fis1* and *mfn1* genes are playing a critical role in erythropoiesis and, therefore, were studied further.

Mitochondrial dynamics and morphology in human erythropoiesis (Fig.1) were studied through *in vitro* differentiation of EPO-induced primary CD34+ hematopoietic stem cells that were isolated from umbilical cord blood and cultured for up to 16 days. Samples were collected at days 0, 3, 5, 8, 10, 13 and 16 of differentiation, pooled, mixed, immunolabeled with anti-CD71-APC and anti-GPA-PE, and cell sorted by for FACS. Four cell populations, representing erythroid progression, were sorted out. R1, progenitors; R2, proerythroblasts; R3, basophilic erythroblasts (hemoglobin is synthetized); and R4+R5, poly and ortho chromatophilic erythroblasts ^29^ (Fig.1A). Then, mitochondria of each cell population were stained with Mitotracker Red for mitochondrial biomass and circularity analysis (Fig. 1B) or with TMRE for mitochondrial membrane potential analysis (Fig. 1 C). Cells were. visualized under confocal microscope. Throughout erythroid differentiation, mitochondrial biomass continuously decreased from R1 (progenitors) to R4+R5 (poly and orthochromatophilic erythroblasts). Circularity, a form factor where 1 is a perfect round mitochondrion and 0 a is filamented or branched mitochondrion, shows that R1 population displayed round mitochondria with a circularity peak value of 0.9. R2 displayed a value of 0.5, meaning that mitochondria became more elongated. Mitochondria circularity moved back to 0.7 in R3 and R4+R5, becoming again fragmented. Mitochondrial membrane potential (Fig 1 C) increased from R1 to R2, and then decreased from R2 to R4+R5. The more elongated mitochondria observed in R2 are concomitant with the highest membrane potential.

**Figure 1.**
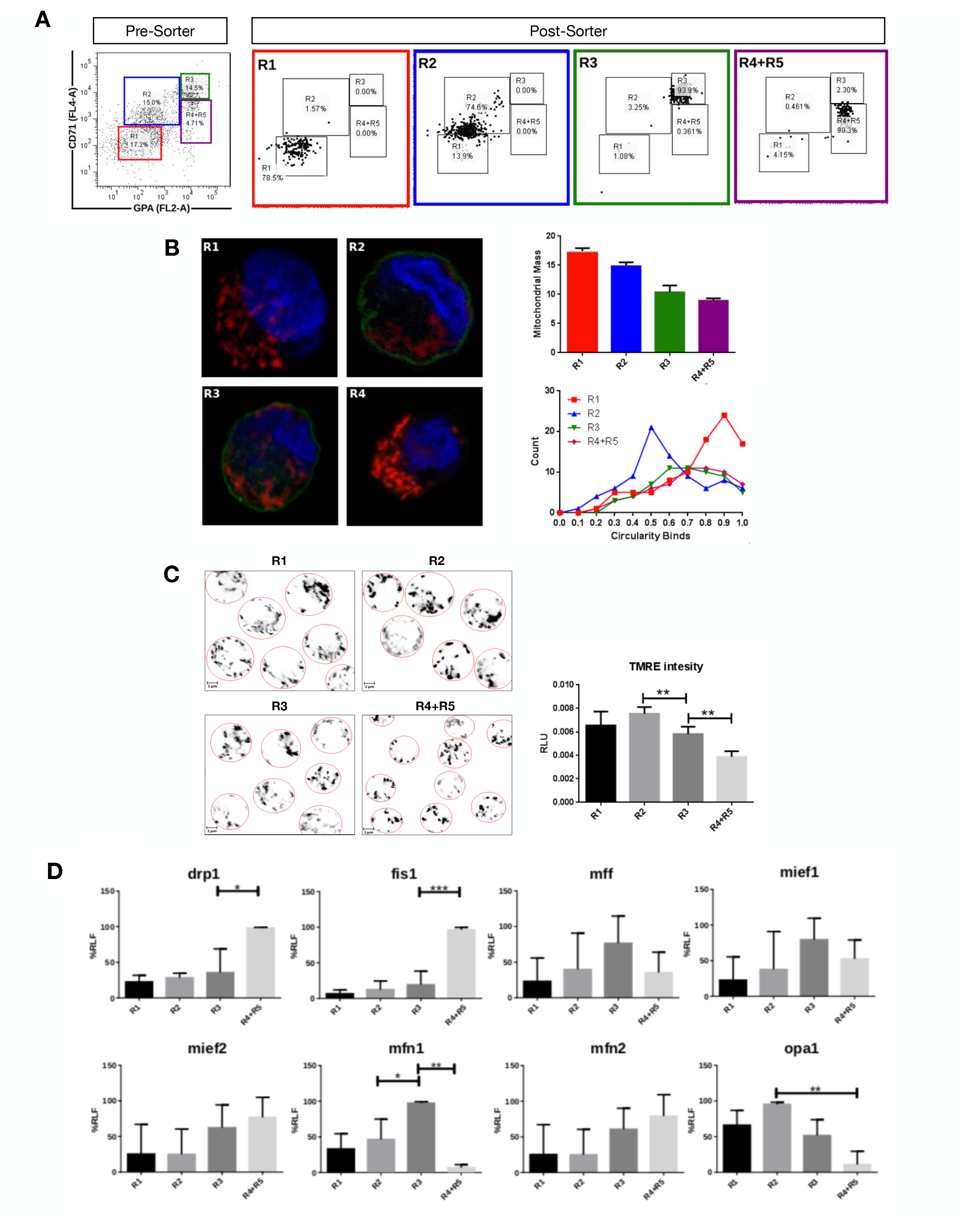
Mitochondrial features throughout erythroid progression. Human hematopoietic CD34+ stem cells isolated from umbilical cord were cultured and EPO-induced to erythropoiesis for 3, 5, 8, 10, 13 and 16 days and pooled together in just one pre-sort sample. **A)** Cell separation of different erythroid populations. Presort samples were labeled with CD71 and GPA surface markers to follow up erythroid progression. Five erythroid populations are distinguished: R1 progenitors, R2 proerythroblast, R3 basophilic erythroblast (these cells start making hemoglobin), R4 polychromatophilic erythroblast and R5 orthochromatophilic erythroblast and reticulocytes. After labelling, samples were sorted to obtain 4 highly pure cell populations as seen as in the post-sort panel for R1, R2, R3 and R4+R5. **B)** Mitochondrial biomass and morphology analysis. Each sorted cell population was stained with DAPI (nucleus, blue) and Mitotracker Red FM (mitochondria, red), and immunolabelled with anti CD71 antibody, green. Confocal microcopy was done to quantify mitochondrial mass and circularity, a form factor where 1 is a perfect circle and 0, an elongated or branched mitochondrion. From R1 to R4+R5, mitochondrial biomass decreases and circularity goes from round mitochondria in R1 to elongated in R2 and then to come back to intermediate values in R3 and R4+R5. **C)** Mitochondrial membrane potential analysis. Sorted cell populations were stained with TMRE dye in a non-quenching mode and visualized under confocal microscope. Membrane potential goes from middle in R1 to high in R2. Then, it starts to decrease toward R4+R5 population. **D)** Mitochondrial Dynamics gene expression analysis. Total RNA, cDNA synthesis and RT-PCR were performed in each sorted population for the mitochondrial fission genes *fis1*, *mff*, *mief1* and *mief2*; and for the mitochondrial fusion genes *mfn1*, *mfn2* and *opa1*. *18s* was used as a loading control. In general, fusion is predominant at early stages and the fission, at later stages of erythroid differentiation. All plotted values are +/− SEMs of *n*=3. Statistical analysis was ANOVA followed by Bonferroni post hoc test (* *p*<0.05, ** *p*<0.01, *** *p*<0.001)

To explore MtDy gene expression, total RNA isolation and cDNA synthesis was performed for R1, R2, R3 and R4+R5 erythroid populations. The expression analysis of the fission genes *fis1, mff, mief 1 and mief* 2; and the fusion genes *mfn1*, *mfn2* and *opa1* was assessed by relative RT-PCR (Fig. 1D). For mitochondrial fission, significant differences were found for *fis1* and *drp1* transcripts, which were upregulated throughout differentiation reaching maximal expression in R4+R5. *Fis1* and *Drp1* had 4 and 1.5 times more transcripts as compared with R3 population (*p*<0.05). On the other hand, for mitochondrial fusion, a significant difference was found for *mfn1* which was upregulated in R3 having twice more transcripts than R2 population. Non-significant differences were observed for *mff*, *mief1*, *mief2, mfn2 and opa1*. These results are in accordance with those obtained in mouse G1E-ER cells and suggest that *fis1* and *mfn1* are important genes for erythroid differentiation. Furthermore, MtDy gene expression profile correlates with circularity values suggesting a predominant mitochondrial fusion from R1 to R2 where mitochondria goes from fragmented to elongated; and then, predominant mitochondrial fission from R2 to R4+R5 where mitochondria became fragmented again. Fragmented mitochondria with low membrane potential in R4+R5 population is imperative for mitochondrial clearance by mitophagy which begins in the orthochromatophilic erythroblasts ^30^

### 3.2 Functional characterization of FIS1

Loss and gain of function experiments for FIS1 were performed by transducing CD34+ cells with lentiviral particles carrying on either pLVCTH-siRNA-FIS1 construct to knock-down FIS1 (FIS1 KD), or pWPI-FIS1 (full cDNA) for FIS1 over-expression (FIS1 OX). It was obtained over 90% efficiency at D3 for both constructs with almost no dead cells (**Fig. S2**). FIS1 KD and FIS1 OX protein expression were confirmed by Western blot which showed a significant 34% reduction and 36% increase respectively in the FIS1 protein levels (*p*<0.01) as compared with controls (**Fig. 2A**).

**Figure 2.**
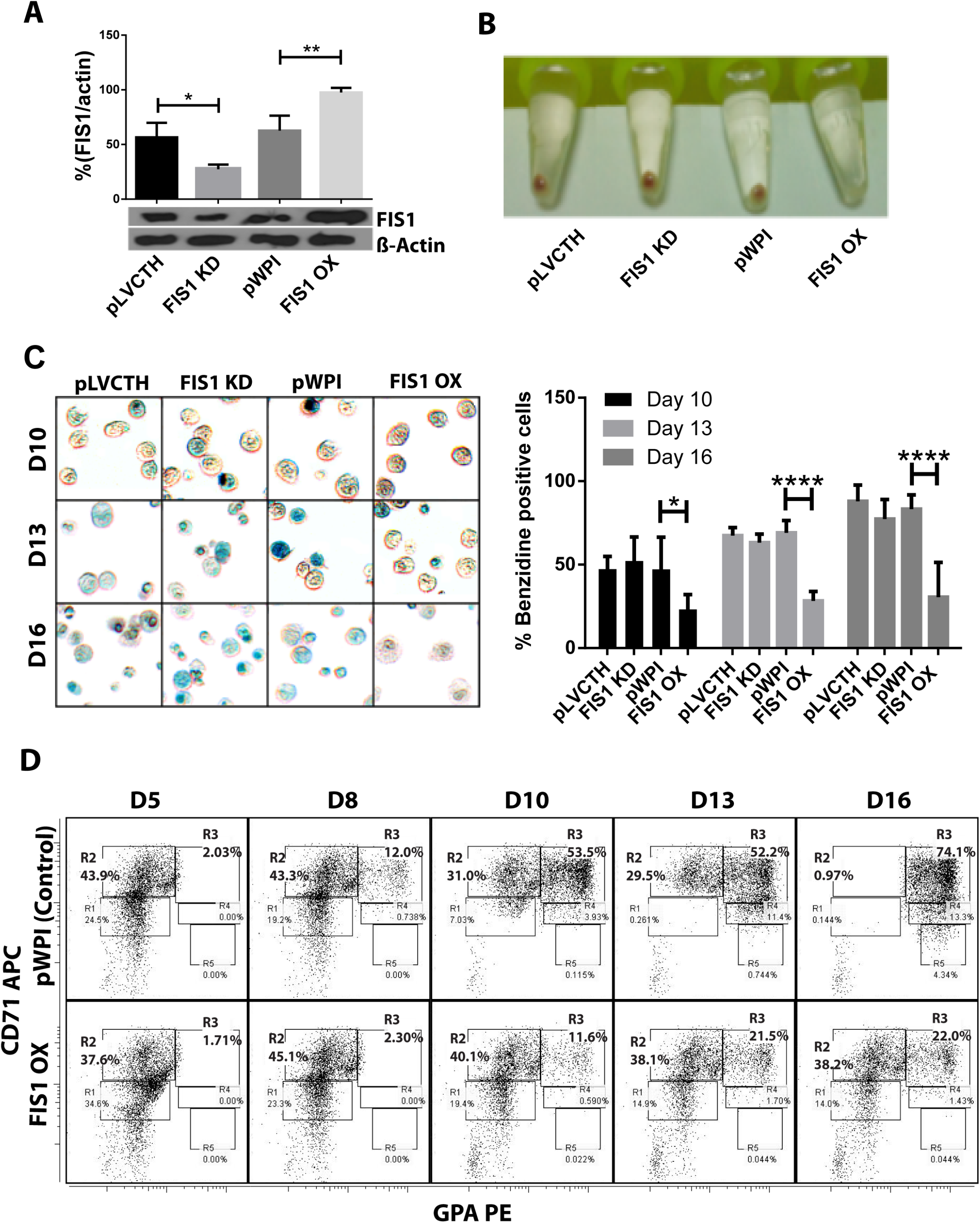
Functional characterization of FIS1 in erythropoiesis. **(A)** Western blot to detect the knock-down or over-expression of FIS1 protein. Quantification was performed by band densitometry. Values are +/− SEM, *n*=3. **(B)** Visual inspection of cellular pellet. FIS1 OX cells do not differentiate under EPO-induced erythroid differentiation. Cell cultures at D16 were spun down and the pellet’s color, visualized. Red pellets were observed in all conditions, but not in FIS1 OX cells whose pellet was white. **(C)** Quantification of hemoglobin carrying cells. Cells from D10, D13 and D16 of differentiation were staining with benzidine to detect hemoglobin. Blue cells (benzidine +) were counted. FIS1 OX cells do not make hemoglobin. Plotted values are +/− SEM, *n*=3. **(D)** Analysis of erythroid progression. FIS OX cells are mainly arrested at the level of progenitors and proerythroblasts. This was seen by flow cytometry analysis of control and FIS OX cells labeled with the surface markers anti-CD71-APC and anti-GPA-PE at D5, D8, D10, D12 and D16 of EPO-induced differentiation. (*n*=3). Statistical analysis, ANOVA followed by Bonferroni post hoc test (* *p*<0.05, ** *p*<0.01, **** *p*<0.001).

Transduced CD34+ cells were EPO-induced to erythroid differentiation for 16 days and spun down to observe their ability to differentiate into red blood cells by looking at color of the pellet, which is expected to be red due to hemoglobin biosynthesis (**Fig. 2B**). While the controls, pLVCTH and pWPI, and the FIS1 KD cells had an intense red pellet, the FIS1 OX cells showed a white cell pellet, suggesting a defect in erythropoiesis. To follow-up hemoglobin biosynthesis, cells were collected at D10, D13 and D16, and benzidine-stained for hemoglobin detection (**Fig. 2C**). Benzidine-positive cells (blue color cells) increased in pLVCTH, pWPI and FIS1 KD conditions reaching over 75% at D16. On the other hand, FIS1 OX cells barely reached 21 % (*p*<0.01).

To understand what stage of erythroid differentiation was affected by FIS1 OX, cells were collected at D5, D8, D10, D13 and D16, and immunophenotyped with the surface markers CD71 and GPA (**Fig. 2D**) for flow cytometry analysis. FIS1 OX cells underwent a delay in erythroid progression starting at D10 with 11.6% of cells in R3 as compared with 53.5% for pPWI control cells. At D13 and D16, FIS1 OX cells displayed 21.5% and 22.0% of R3 cells respectively as compared with 52.2% and 74.1 % of R3 cells for control cells. Besides, at D16, FIS1 OX cells displayed 38.2% R2 cells as compared with 0.97% R2 cells for control cells. R3 population is featured by the onset of hemoglobin biosynthesis. On the other hand, FIS1 KD cells showed a normal differentiation pattern as compared with pLVCTH control cells **(Fig. S3)**. These results suggest that exacerbation of mitochondrial fission at early stages of erythroid differentiation disrupts hemoglobin biosynthesis and delays erythroid progression.

### 3.3 Effects on mitochondrial parameters

Assessment of mitochondrial morphology was performed in FIS1 OX cells (**Fig. 3**) and FIS1 KD cells **(Fig. S4)** from D5 to D16 of differentiation by immuno-staining with anti-VDAC1 antibody (red channel). Mid-sagittal optical sections of positive-transduced cells (GFP+) were analyzed by confocal microscopy. As expected, FIS1 OX cells displayed a fragmented mitochondrial web with some large and round mitochondria as compared with pWPI control cells (**Fig. 3A**). Mitochondrial morphology analysis in term of circularity, area and perimeter revealed a very well-structured morphological pattern during erythroid differentiation in control cells. A positive correlation (positive slope) was found between circularity and area (**Fig. 3B**); and between circularity and perimeter (**Fig. 3C**) with a correlation coefficient (R^2^) of 0.78 - 0.87 and 0.88 – 0.91, respectively. Mitochondrial area (**Fig. 3B**) and perimeter (**Fig. 3C**) distribution (seen on the right-Y axis) shifted slightly toward smaller values during differentiation and circularity distribution (seen on the upper-x axis) ranged from 0 to 0.5 having a peak of 0.2 in control cells. On the other hand, the overexpression of FIS1 in FIS1 OX cells caused major changes in mitochondrial morphology which were evident as early as D5 and throughout erythroid differentiation as compared with control cells. Firstly, the correlation between circularity and area is lost with R^2^ values of 0.01 to 0.26 mainly because circularity distribution shifted considerable toward higher values (ranging from 0.6 to 1 with a peak of 0.9) with mitochondria having a larger area than control cells (**Fig. 3B**). Furthermore, a negative correlation (negative slope) was found between circularity and perimeter with a R^2^ of 0.93; and the perimeter switched from a normal to a bimodal distribution from D10 (**Fig. 3C**). These quantitative analyses of mitochondrial morphology for FIS1 OX cells are in agreement with the presence of a fragmented mitochondrial web with a greater amount of large and round mitochondria and suggest that unbalanced MtDy favoring mitochondrial fission are responsible for disruption of both heme biosynthesis and erythroid progression. Regarding FIS1 KD cells, no significant differences were found as compared with controls. (**Fig. S4**).

**Figure 3.**
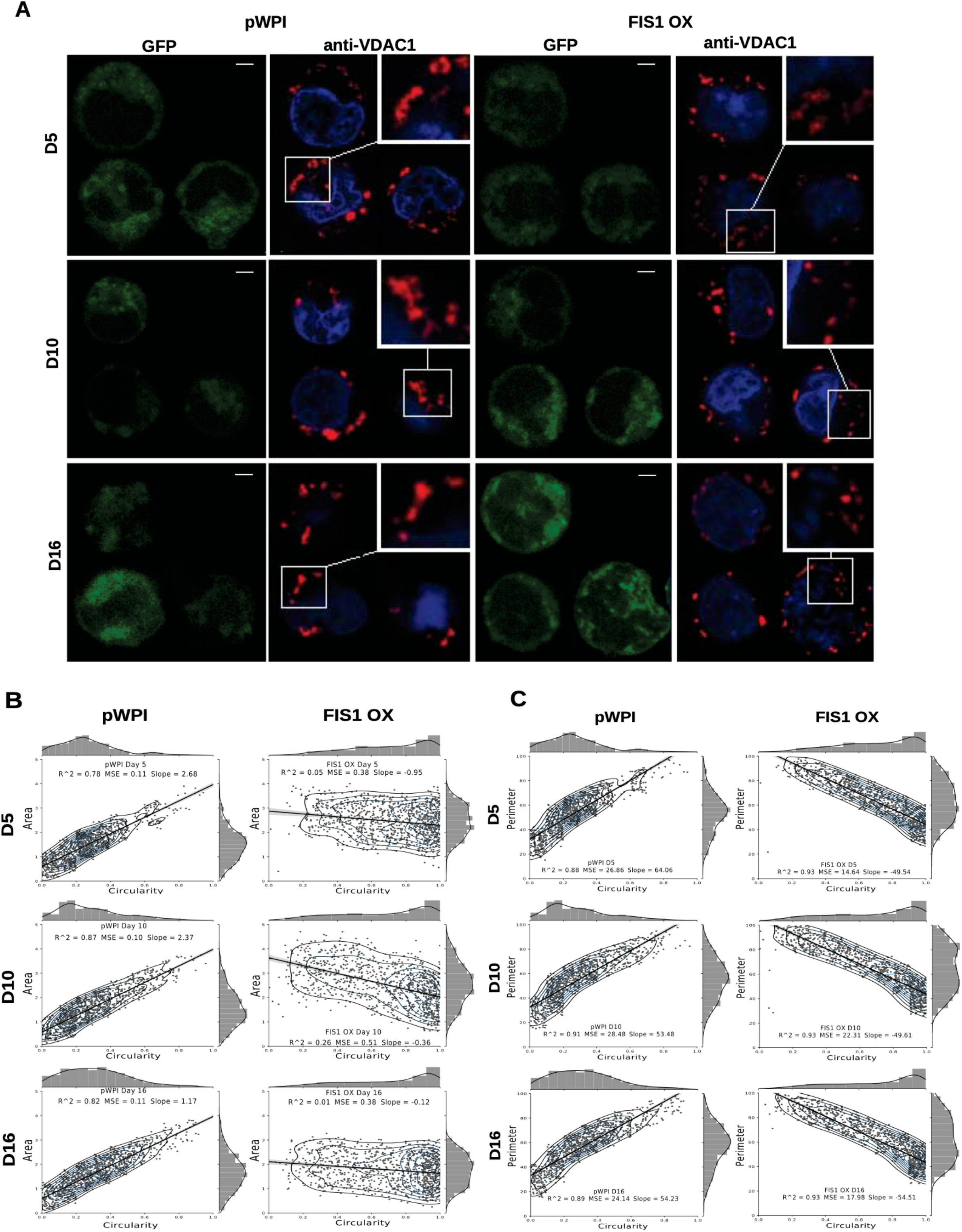
Effects of FIS1 OX on mitochondrial morphology. **(A)** Mitochondria visualization by confocal microscopy. Immunofluorescence microscopy at D5, D10 and D16 of erythroid differentiation for pWPI control and FIS1 OX cells. Anti-VDAC1 antibody labeled mitochondria (red) and the DAPI dye, the nucleus (blue). **(B)** Mitochondrial morphometric analysis in term of Area at D5, D10 and D16 of differentiation for control pWPI and FIS1 OX cells. It was performed using the Z-slides from GFP+ cells. Each dot represents a mitochondrial unit in terms of Area (Y axes) and Circularity (X axes). Also, a frequency histogram of mitochondrial circularity and area were added. Linear regression analysis was performed to compare slopes., which were significantly different (*p*<0.05). R^2^ and MSE were also calculated **(C)** Similar to (B). Mitochondrial morphometric analysis in term of Perimeter. Each dot represents a mitochondrial unit in terms of Perimeter (Y axes) and circularity (X axes).

To gain insights into the bioenergetic capacity of mitochondria in FIS1 OX cells, the OXPHOS protein expression and the mitochondrial membrane potential were assessed at day 10, 13 and 16 (**Fig. 4**). At the level of protein expression, complex II significantly decreased by 50% (*p*<0.01) and complex IV by 46% (*p*<0.05) as compared with control pWPI cells (**Fig. 4A**). Similarly, mitochondrial membrane potential decreased 20% in FIS1 OX cells at day 10 (*p*<0.001) and 16 (*p*<0.05) (**Fig. 4B**). These results suggest that disruption of MtDy by FIS1 over-expression decreases the bioenergetic capacity of mitochondria.

**Figure 4.**
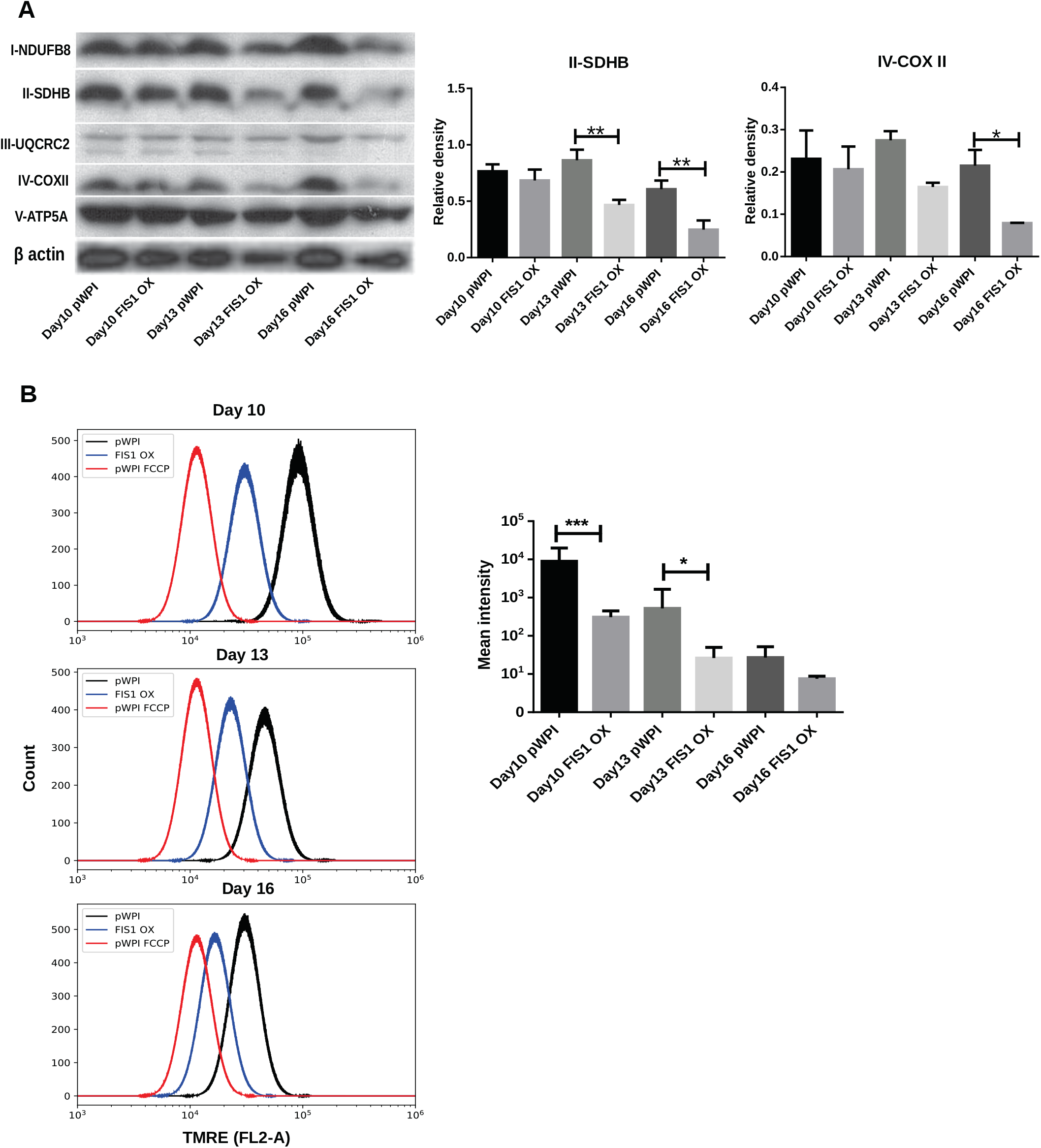
Effects of FIS1 OX on mitochondrial bioenergetics. **(A)** OXPHOS protein expression levels. Total protein lysate from FIS1 OX and pWPI control cells at D10, D13 and D16 of erythroid differentiation were assayed for immunoblot to detect the OXPHOS protein expression level. Band densitometry analysis was performed for Complex II and Complex IV. The plotted values are +/− SEM. *n*=3. Statistical analysis, ANOVA followed by Bonferroni post hoc test (* *p*<0.05, ** *p*<0.01) **(B)** Measurement of mitochondrial membrane potential. FIS1 OX and pWPI control cells at D10, D13 and D16 erythroid differentiation were stained with 10nM TMRE in non-quenching mode and analyzed by flow cytometry. 10uM FCCP was used as negative control. TMRE mean intensity was quantified. The values plotted are +/− SEMs. *n*=3. Statistical analysis, ANOVA followed by Bonferroni post hoc test (* *p*<0.05)

### 3.4 Effect of FIS1 OX on cell survival during erythroid differentiation

Our results have shown a loss of mitochondrial homeostasis which may lead to programmed cell death. We asked if apoptosis was responsible for the disruption in hemoglobin biosynthesis and delayed erythropoiesis in FIS1 OX cells. By Western blot, the levels of pro-apoptotic active caspase 9 (intrinsic pathway), active caspase 8 (extrinsic pathway), and active caspase 3 (apoptosis executioner) were measured in differentiating FIS1 OX and control cells at D10, D13 and D16 (**Fig. 5**). No significant differences were found in cleaved caspase 9 (**Fig. 5A**) and caspase 8 (**Fig. 5B**). While caspase 3 showed an initial difference at Day 10 in FIS1 OX, this was not sustained throughout following days (**Fig. 5C**). Caspases have been reported to play a role in erythropoiesis, in particular caspase 3 ^31^ and that may explain the presence of active proteins. However, since no differences were observed between FIS1 OX and control cells, these results suggest that apoptosis is not triggered by FIS1 overexpression and therefore it is not a factor involved in the hemoglobin and erythropoiesis disruption seen in FIS1 OX cells.

**Figure 5.**
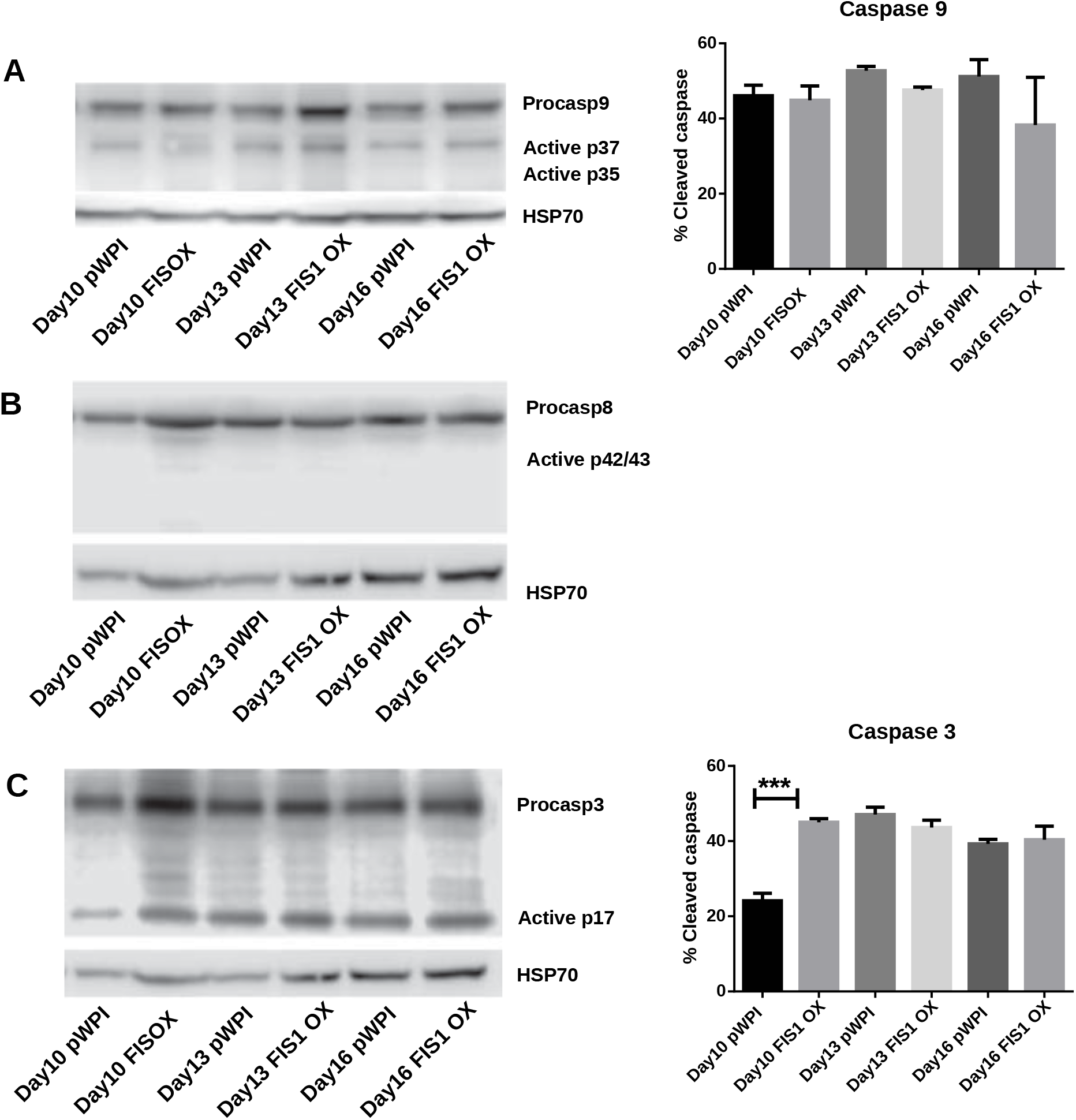
Assessment of apoptosis in FIS1 OX cells. Total protein lysates from FIS1 OX and pWPI control cells at D8, D10, D13 and D16 of erythroid differentiation were examined by Western blot. **(A)** Protein levels of Procaspase 9 and cleaved active forms P37 and P35. This is for intrinsic apoptosis. HSP70 were used as housekeeping control. Band densitometry for p37, as a percentage of total caspase 9, was plotted. Values are +/− SEM. *n*=3. Statistical analysis, ANOVA followed by Bonferroni post hoc test. **(B)** Protein levels of Procaspase 8 and cleaved active forms P42/P43. This is for extrinsic apoptosis. HSP70 were used as housekeeping control. **(C)** Protein levels of Procaspase 3 and cleaved active form P17. This is a general apoptotic effector. HSP70 were used as housekeeping control. Band densitometry for p17, as a percentage of total caspase 3, was plotted. Values are +/− SEM. *n*=3. Statistical analysis, ANOVA followed by Bonferroni post hoc test (*** *p*<0.001).

### 3.5. MFN1 knock-down (MFN1 KD) delays erythroid differentiation

To confirm whether a MtDy is a main actor in the erythropoietic cell differentiation, MFN1 was knocked down to mimic the FIS1 OX phenotype i.e. the fragmentation of the mitochondrial web and the appearance of large and round mitochondria. It has been reported that MFN1 KD triggers round mitochondria morphology in several cells models ^32–34^. Right after isolation, CD34+ cells were transduced with pLVCTM-shMFN1 lentiviral particles to knock-down MFN1 protein. MFN1 depletion was confirmed by Western blot, which showed a significant 58% reduction in MFN1 protein levels (*p*<0.001) (**Fig. 6A**). Mitochondria-morphological analysis of MFN1 KD cells revealed a distinct pattern as compared with FIS1 OX cells. MFN1 KD cells displayed a fragmented but homogeneous mitochondria population in terms of area, perimeter and circularity during erythroid differentiation (**Fig. 6**) as compared with FIS1 OX cells (**Fig. 3**). MFN1KD mitochondria exhibited mainly a reduction in size and an increase in circularity i.e. mitochondria became rounded and smaller than control cells. Comparing FIS1 OX with MFN1 KD mitochondria population, the former is an average of 3 times bigger than the latter (**Fig. 6C** versus **Fig. 3B**). Erythroid progression analysis by flow cytometry showed that MFN1 KD caused a delay in erythroid progression but did not disrupt heme biosynthesis. This was evident at day 16 looking at R2 and R3 populations. R2 cells are 14.3% and R3, 63.7% in MFN1 KD cells versus 4.14% and 73.7% respectively in controls cells. (**Fig. 6E**).

**Figure 6.**
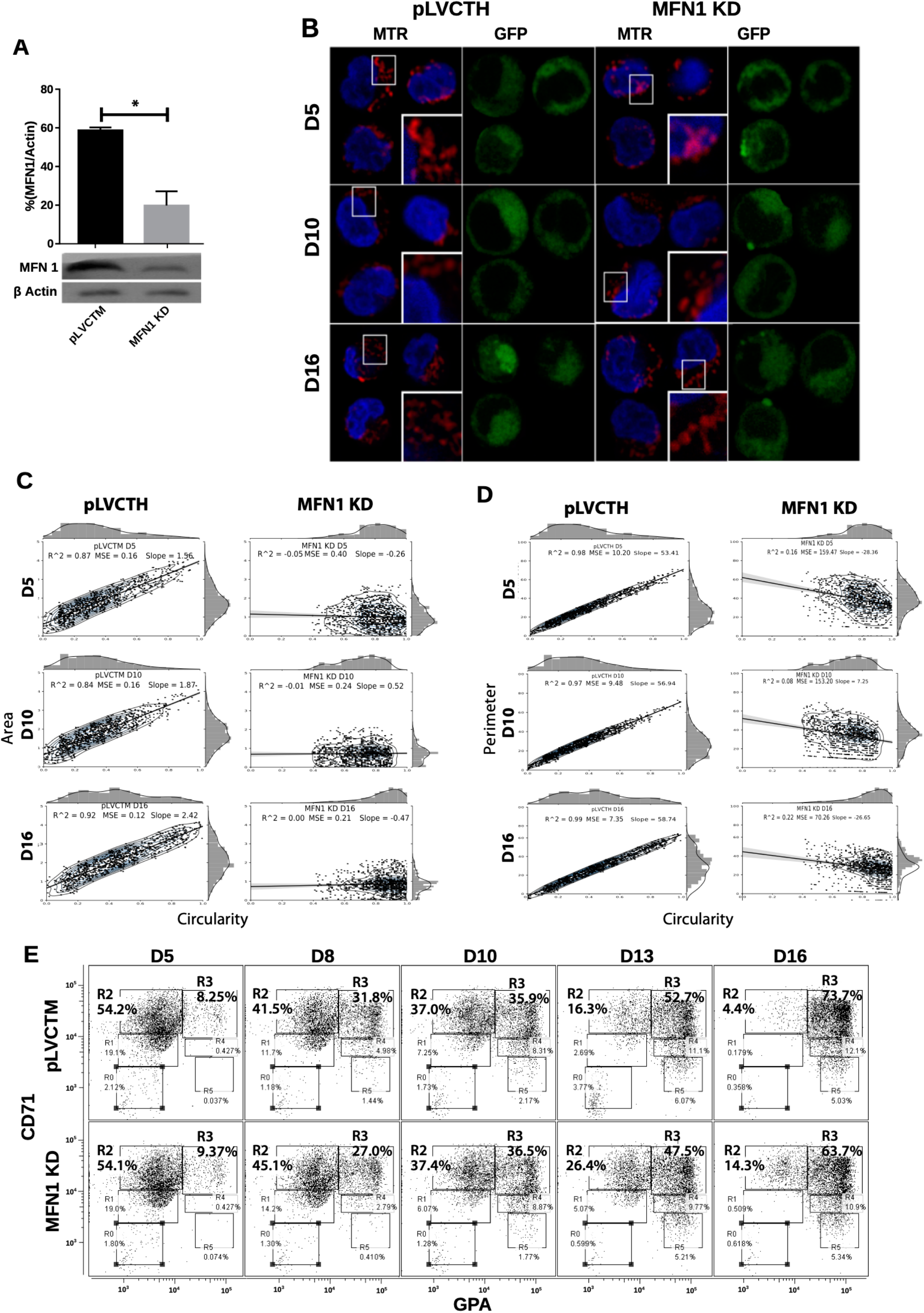
Effect of MFN1 KD on mitochondrial morphology and erythroid differentiation. **(A)** Western blot to detect the knock-down of MFN1. Total protein lysates from pLVCTH control and MFN1 KD cells at D3 of erythroid differentiation were examined by Western blot to check MFN1 expression. Quantification was performed by band densitometry. Plotted values are +/− SEM. *n*=3. Statistical analysis, ANOVA followed by Bonferroni post hoc test (* *p*<0.05). **(B)** Mitochondria visualization by confocal microscopy. Immunofluorescence microscopy at D5, D8, D10, D13 and D16 of erythroid differentiation for pLVCTH control and MFN1 KD cells. Anti-VDAC1 (red) labeled mitochondria and DAPI (blue), the nucleus. **(C)** Mitochondrial morphometric analysis in term of Area at D5, D10 and D16 of erythroid differentiation for pLVCTH control and MFN1 KD cells. It was performed using the Z-slides from GFP+ cells. Each dot represents a mitochondrial unit in terms of area (Y axes) and circularity (X axes). Also, a frequency histogram of mitochondrial circularity and area were added. Linear regression analysis was performed to compare slopes, which were significantly different (*p*<0.05). R^2^ and MSE were also calculated. **(D)** Similar to (C). Mitochondrial morphometric analysis in term of Perimeter. Each dot represents a mitochondrial unit in terms of Perimeter (Y axes) and circularity (X axes). **(E)** Erythroid progression in MFN1 KD cells. Flow cytometry analysis of surface markers anti-CD71-APC and anti-GPA-PE in pLVCTH control and MFN1 KD cells at D5, D10 and D16 days of EPO-induced differentiation. All analysis (*n*=3) correspond GFP+ cells. MFN1 KD cells have a bigger R2 and smaller R3 population at D13 and D16 than control cells, meaning a delay in erythroid progression as compared with control cells.

Even though FIS1 overexpression and MFN1 knockdown resulted in mitochondrial web fragmentation, the final output of mitochondrial phenotype was distinctive and associated with divergent physiological consequences in erythropoiesis. These interesting results suggest that the larger mitochondrial size in FIS1 OX mitochondria might be given by other mechanism coupled to MtDy through the action of FIS1 protein.

### 3.6. Closure of the mitochondrial permeability transition pore (mPTP) rescues the erythroid differentiation in FIS1 OX cells

The appearance of large and round mitochondria with low membrane potential and reduced OXPHOS expression, along with the disruption of heme biosynthesis and the halt of erythropoiesis in FIS1 OX cells, suggested the involvement of the mitochondrial permeability transition pore (mPTP). Transmission electron microscopy (TEM) was used to analyze mitochondrial ultrastructure and provide further insight into this process **(Fig. 7A)**. Control cells displayed mitochondria with normal morphology, matrix density and cristae structure at D13 and D16 of differentiation. On the other hand, FIS1 OX cells displayed a heterogeneous mitochondria population with immature-like mitochondria with few a poor developed cristae and swollen-like mitochondria (**Fig 7A**). These ultrastructure images of mitochondria, which are in agreement with the morphology results obtained by confocal microscopy (Fig.3), suggested the involvement of the mPTP as a potential mechanism of signaling between MtDy and the erythropoietic phenotype. To corroborate this, FIS1 OX and control cells were treated with the mPTP inhibitor cyclosporin A (CsA). TEM images corroborated that CsA treatment rescued mitochondrial morphology. CsA-treated FIS1 OX cells had mitochondria with normal size and cristae structure similar to control cells (**Fig. 7B)**; and the quantitative analysis of mitochondrial circularity (**Fig. 7C**) confirmed the recovery in CsA-treated FIS1 OX cells. Finally, flow cytometry analysis of cells collected at D10, D13 and D16 of erythroid differentiation showed no differences between CsA-treated control cells and CsA-treated FIS OX; and that hemoglobin biosynthesis was also rescued (**Fig.7D**). Ours findings suggest that changes in MtDy towards a more fragmented mitochondrial web due to FIS1 over expression correlates with the opening of the mPTP, which in turns arrested erythroid differentiation and inhibited hemoglobin biosynthesis.

**Figure 7.**
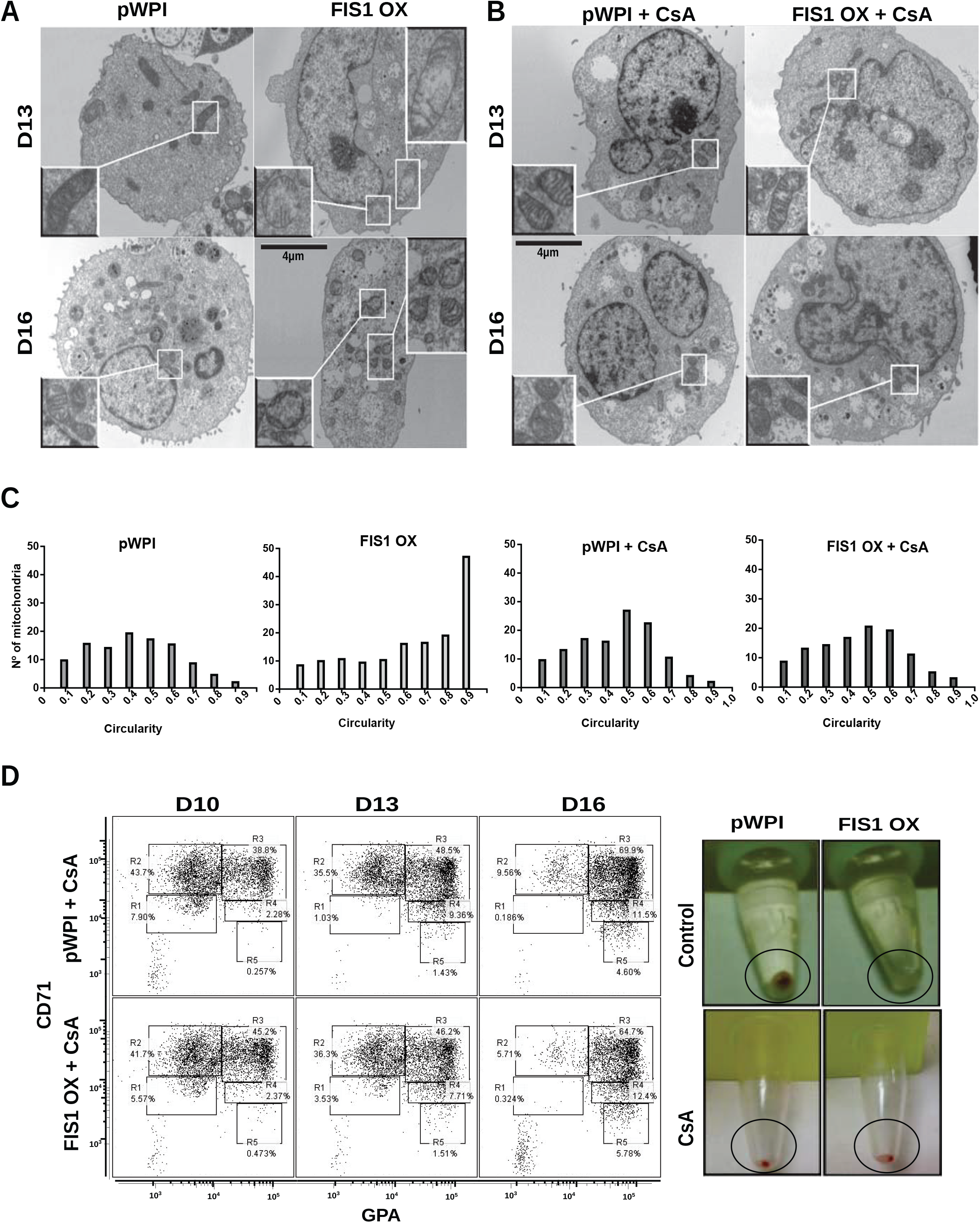
Assessment of the mPTP in FIS1 OX cells. **(A)** Ultrastructure visualization of mitochondria. Transmission electron microscopy (TEM) was performed for pWPI control and FIS1 OX cells at D13 and D16 of erythroid differentiation. FIS1 OX displayed heterogeneous mitochondrial morphologies. Some mitochondria look like immature mitochondria with short and few cristae. It was also found swollen-like mitochondria as compared with control cells. Magnification 6000x. **(B)** Ultrastructure visualization of mitochondria in CsA-treated cells. TEM analysis showed normal mitochondrial structure and electron density in CsA-treated FIS1 OX cells as compare with CsA-treated pWPI control cells. **(C)** Mitochondrial circularity analysis. It was calculated from TEM images of pWPI control and FIS1 OX cells treated or not treated with CsA at D13 of erythroid differentiation. Histograms showed that the treatment with CsA fully rescued the mitochondrial morphology. **(D)** Erythroid progression analysis of CsA-treated FIS1 OX cells. Cells were treated with CsA on D9 and collected at D10, D13 and D16. Right after collection, cells were immunolabeled with anti-CD71-APC and anti-GPA-PE and analyzed by flow cytometry All analysis (*n*=3) correspond to GFP+ cells. In addition, cells were spun down to observe the cell pellet’s color. Treatment with CsA rescued erythroid differentiation and hemoglobin biosynthesis in FIS1 OX cells.

## 4. Discussion

Fine regulation of mitochondrial fission and fusion is required for energy demands adjustments, metabolic control and signaling, allowing proper cellular function ^35^ cell proliferation and differentiation ^36,37^, stem cell commitment ^19^ and cell death. In this work, we contribute to the understanding of MtDy during erythropoiesis. We found that the most expressed fission genes are *fis1* and *drp1*; and the fusion one, *mfn1*. All of them have a well-defined expression pattern, increasing their mRNA levels as erythroid differentiation moves forward. Furthermore, we showed that exacerbated fission disrupted mitochondrial morphology and function, heme biosynthesis and delayed erythroid differentiation, keeping cells in an undifferentiated state; and that the MtDy signaling mechanism involved the mPTP.

The role of FIS1 as a direct receptor of DRP1 to induce mitochondrial fission has been controversial in mammals, because recruitment of DRP1 to mitochondria or DRP1-mediated fission has been shown to be independent of FIS1 expression levels ^8,38,39^. In fact, it was recently published that FIS1 is an inhibitor of mitochondrial fusion, interacting directly with MFN1, MFN2 and OPA1^40^. Thus, mitochondria fragmentation in FIS1 overexpressing cells would be due to the inhibition of mitochondrial fusion. However, other recent papers have shown the efficacy of a molecule able to inhibit the interaction DRP1/FIS1, which blocks pathological or excessive mitochondrial fragmentation seen in ALS and cardiac diseases without affecting basal mitochondrial fission ^41^. Our results showed that both FIS1 OX and MFN1 KD erythropoietic cells had a delay in erythropoiesis. Nevertheless, this delayed was more evident and stronger in FIS1 OX cells; and only FIS1 OX cells displayed a disruption in heme biosynthesis. Furthermore, the mitochondrial web fragmentation caused by FIS1 OX and MFN1 KD are not alike as revealed by a deep mitochondrial morphological analysis. FIS1 OX cells had a fragmented mitochondrial network but with heterogeneous and bigger mitochondria (large and round mitochondria) than MNF1 KD cells, which also have fragmented mitochondria, but they are homogeneous and smaller. Our data suggest that both FIS1 OX and MFN1 KD are able to cause a mitochondrial web fragmentation, they do not have a redundant role in mitochondrial function in erythropoiesis.

Excessive mitochondrial fission has been associated with a decreased mitochondrial function and increased ROS ^41,42^. FIS1 up-regulation decreased cellular ATP levels in anoxic cardiomyocytes and impaired glucose-stimulated insulin secretion in INS-1E cells ^43,44^; and it has been identified as a molecular marker for poor prognosis in patients with acute myeloid leukemia^45^. We observed that erythroid differentiation under FIS1 overexpression was arrested mainly at the proerythroblast level with mitochondria featuring reduced complex II and IV protein expression and membrane potential; and an immature phenotype i.e. short and poor developed cristae as seen by TEM. No dead or apoptotic cells were observed, which suggested that FIS1 overexpression induced mitochondrial metabolic reprogramming that maintains cells in the glycolytic state rather than shifting to oxidative metabolism to support cell proliferation. Furthermore, our results supports the evidence of excessive levels of FIS 1 as a molecular marker of myeloid leukemia ^45^.

As we discussed previously, disruption of mitochondrial fusion by MFN1 KD delays erythroid progression. Several studies about cell reprogramming have shown that depletion of MFN1 inhibit the p53-p21 pathway, promoting the conversion of somatic cells to pluripotent cells ^46^. Also, depletion of MFN1 in mouse embryonic heart cells stalled the heart development through impairments in the cell differentiation to cardiomyocytes ^47–49^. The role of MtDy in commitment and differentiation of stem cell has been also shown in neural stem cells for neurogenesis ^50,51^These evidences strengthen our results that changing the balance of mitochondrial dynamics towards a more fragmented state is able to induce a metabolic reprogramming affecting the differentiation process.

Although the role of MtDy for cell physiology is well accepted, there is not clarity about the signaling mechanisms to induce the metabolic reprogramming which involves transcriptional regulation. Our results suggest that the mitochondrial permeability transition pore, mPTP, is a link between MtDy and the signaling mechanisms to control cell differentiation. FIS1 OX generated large and round mitochondria, which were not seen in MFN1 KD cells. Treatments with cyclosporin A (CsA), which is an inhibitor of the mPTP, rescued the mitochondrial morphology, the heme biosynthesis and the progression of erythropoiesis. Interestingly, the role of the mPTP in cellular differentiation has been reported in the past for early embryonic cardiomyocytes, where a frequent opening of the mPTP is needed to maintain an immature mitochondrial morphology with low oxidative capacity. On the other hand, the closure of the pore was required for proper differentiation into cardiac muscle cells ^52,53^. Furthermore, mPTP flashes are needed for cortical neural progenitor differentiation ^54^; and closure of mPTP with CsA is protective against extra physiologic oxygen shock/stress in hematopoietic stem cell transplantation ^55^; and needed for a more efficient *in vitro* differentiation of CD34+ cells to red blood cells ^56^.

One of the triggers of mPTP opening is an elevated Ca^++^ concentration in the mitochondrial matrix ^57^. In this regard, FIS1 can interact with the ER protein BAP31 allowing mitochondria to be loaded with Ca^++^ ^58,59^ and then establishing a potential mechanism between mitochondrial dynamics and the mPTP.

In conclusion, the proper balance of MtDy is essential for erythroid differentiation. Disruption of MtDy by FIS1 overexpression in hematopoietic stem cells altered mitochondrial function, delayed their differentiation into red blood cells and blocked heme biosynthesis, keeping them in an immature and proliferative state. The red blood cell differentiation in FIS1 OX cells was fully rescued by the addition of CsA, a pharmacological inhibitor of the mPTP, into the culture media. We have also shown that MFN1 KD caused mitochondrial fragmentation and delayed erythropoiesis, but did not caused the same effects on mitochondrial morphology as FIS OX and did not block heme biosynthesis. Thus, MFN1 and FIS1 are not redundant in their functions. Our results suggest that MtDy is regulating the aperture/closure of the mPTP which in turn will allow/block the exit of signal molecules such a Ca^++^, ROS and other metabolites to manage the genetic and metabolic reprograming of cells for proper cellular differentiation.

## 5. Acknowledgment

This work was funded by the grants: Fondecyt 1100995 (AAE), Fondecyt 1180983 (AAE); DI-209-12/N (AAE); Millennium Institute on immunology and Immunotherapy P09-016-F (AAE) and Ph.D. CONICYT Scholarship 2120552 DN2012 (AG). We also thank the Maternity Service at Complejo Asistencial Barros Luco, Santiago, Chile for its participation in this study by providing us the umbilical cord blood for CD34+ cell isolation.

## 6. Authorship Contribution

Alvaro M. Gonzalez performed most of the experiments and data analysis and helped with manuscript writing. Lina M. Ruiz and Erik Jensen performed TEM analyses. Cesar A. Echeverria and Valentina Romero performed apoptosis experiments. Linsey Stiles and Orian Shirihai designed and performed the qPCR on G1E-ER cells. Alvaro A. Elorza made the experimental design, analyzed data, wrote the manuscript and funded the research.

