## Supplemental Figure Legends for "Exacerbation of mitochondrial fission in human CD34^+^ cells halts erythropoiesis and hemoglobin biosynthesis"

**Supplemental Figure 1. Mitochondrial dynamics in mouse erythropoiesis.**

**(A)** Visualization of mitochondrial morphology. The mouse erythropoietic cell line G1E-ER were ß-estradiol-treated to induced erythroid differentiation. Mitochondria were stained with 10nM MitoTracker Red and visualized by confocal microscopy at 24 hrs. post induction. The mitochondrial web became fragmented upon differentiation.

**(B)** Mitochondrial Dynamics gene expression. mRNA expression kinetics of mitochondrial dynamics genes during ß-estradiol-induced erythropoiesis in G1E-ER cells was assessed by quantitative real time (qRT-PCR). Cells were collected at 0, 12, 24, 36 and 48 hrs.

**Supplemental Figure 2. Lentiviral transduction of CD34+ cells**

CD34+ cells were lentivirus-transduced one hour after isolation overnight. Three days later, transduction efficiency was assessed by flow cytometry. GFP expression was used as indicator of successful transduction. In addition, cells were stained with 7-AAD to assess dead cells. Hydrogen Peroxide was used in control cells as a positive control of dead cells. Transduction efficiency of CD34+ was over 90% with no dead cells (<1%).

**Supplemental Figure 3. Erythroid Progression in FIS1 KD cells**

Flow cytometry analysis of surface markers anti-CD71-APC and anti-GPA-PE in pLVCTH control and FIS1 KD cells at D5, D8, D10, D12 and D16 of EPO-induced erythroid differentiation. (*n*=3).

**Supplemental Figure 4. Effects of FIS1 KD on mitochondrial morphology**

**(A)** Mitochondria visualization by confocal microscopy. Immunofluorescence microscopy at D5, D10 and D16 of erythroid differentiation for pLVCTH control and FIS1 KD cells. Anti-VDAC1 antibody labeled mitochondria (red) and the DAPI dye, the nucleus (blue).

**(B)** Mitochondrial morphometric analysis in term of Area at D5, D10 and D16 of differentiation for control pWPI and FIS1 KD cells. It was performed using the Z-slides from GFP+ cells. Each dot represents a mitochondrial unit in terms of Area (Y axes) and Circularity (X axes). Also, a frequency histogram of mitochondrial circularity and area were added. Linear regression analysis was performed to compare slopes., which were significantly different (*p*<0.05). R^2^ and MSE were also calculated.
