## Supplementary figures and images for "Exacerbation of mitochondrial fission in human CD34^+^ cells halts erythropoiesis and hemoglobin biosynthesis"

### Supplemental Figure 1

##### Supplementary 1

**A**

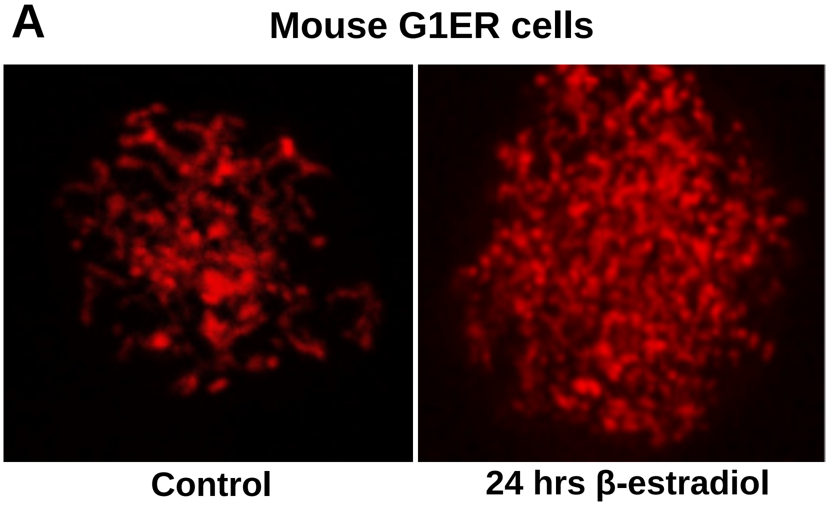


**B**

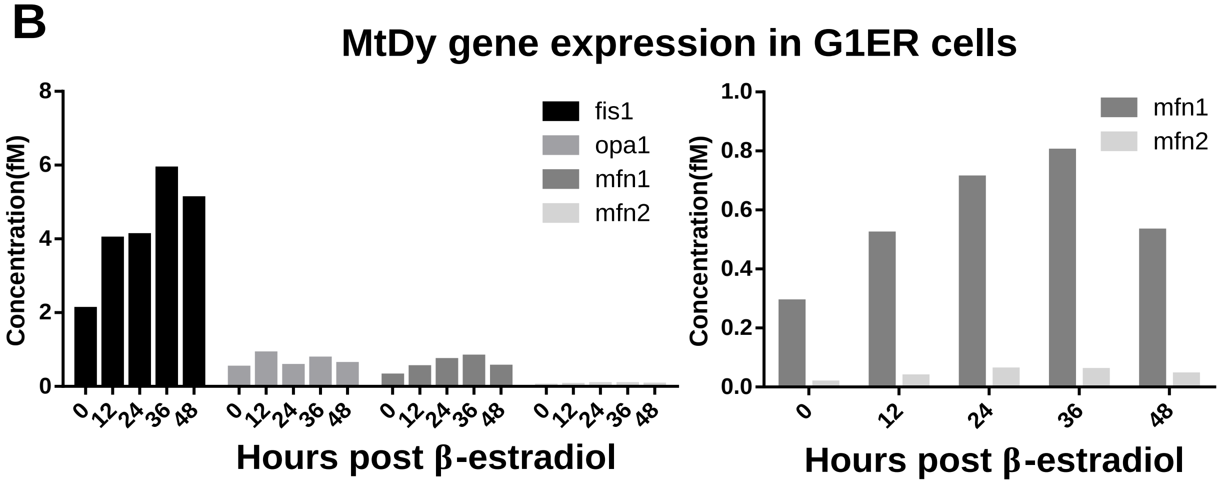

### Supplemental Figure 2

##### Supplementary 2

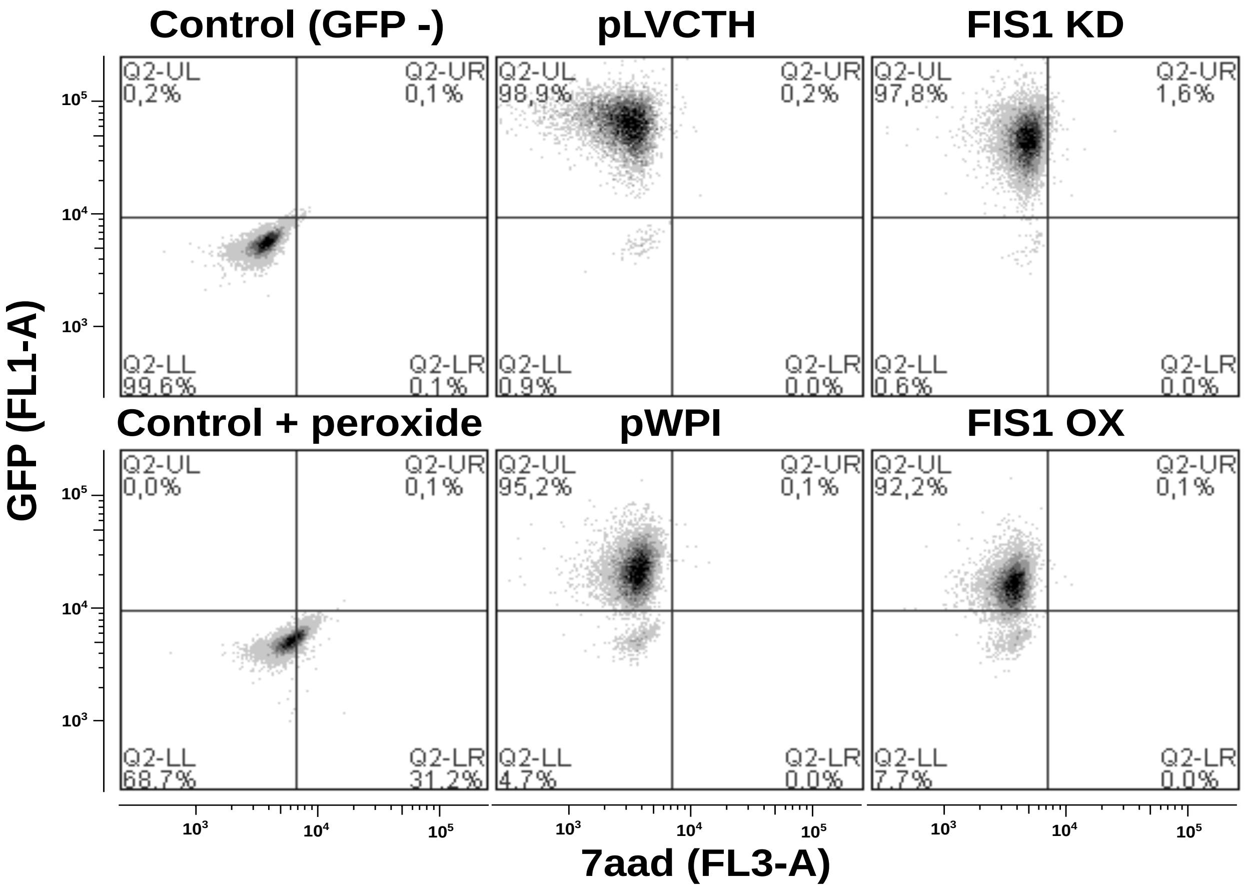

### Supplemental Figure 3

#### Supplementary 3

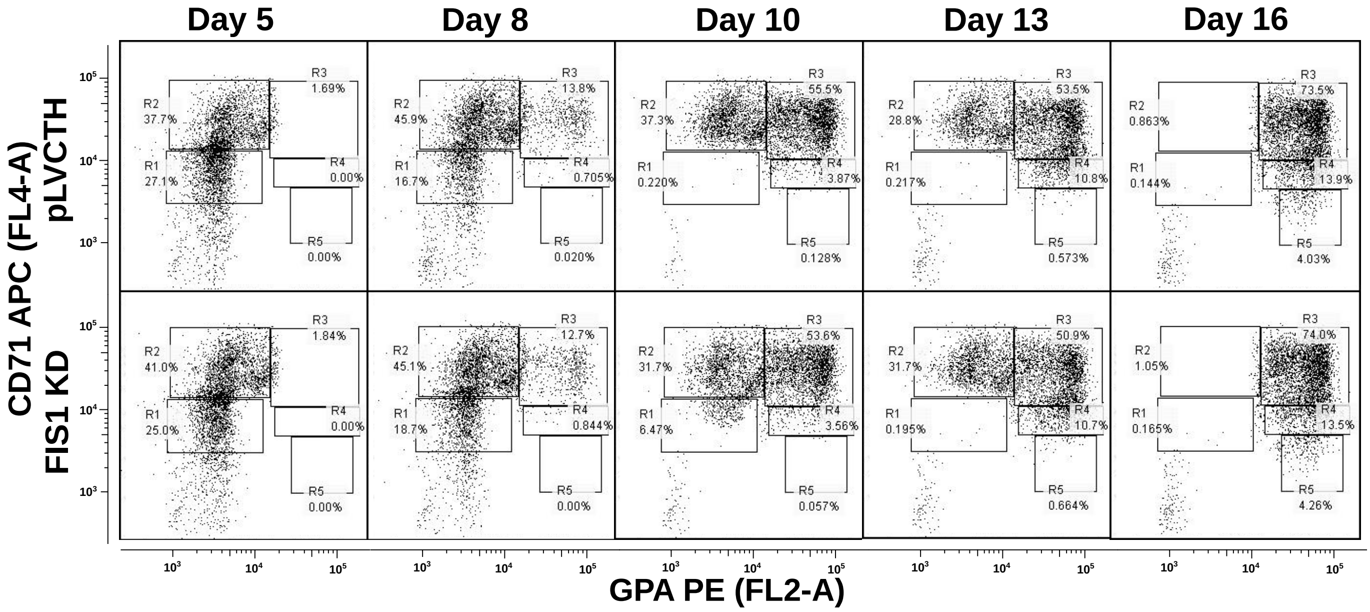

### Supplemental Figure 4

#### Supplementary 4

A

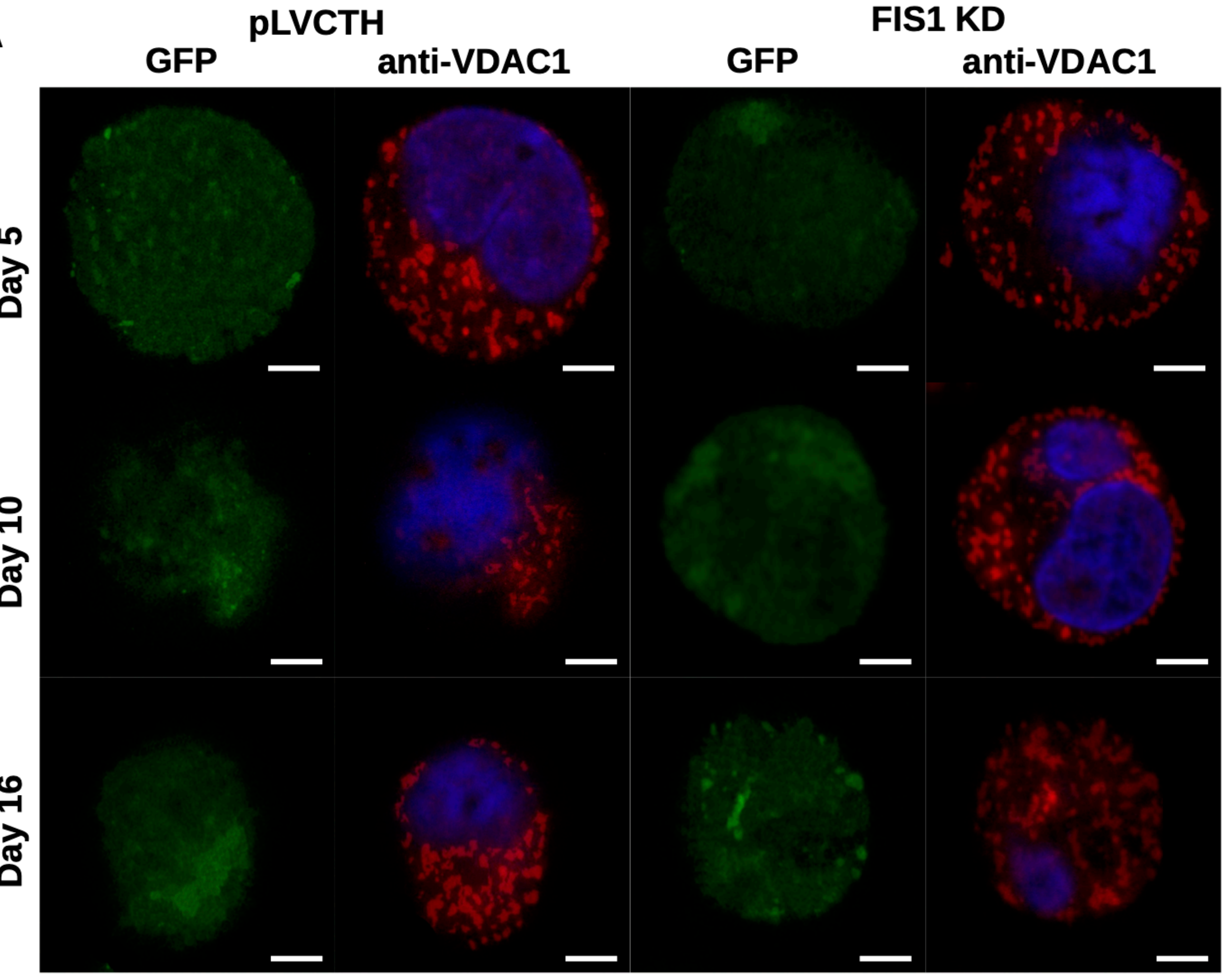


B

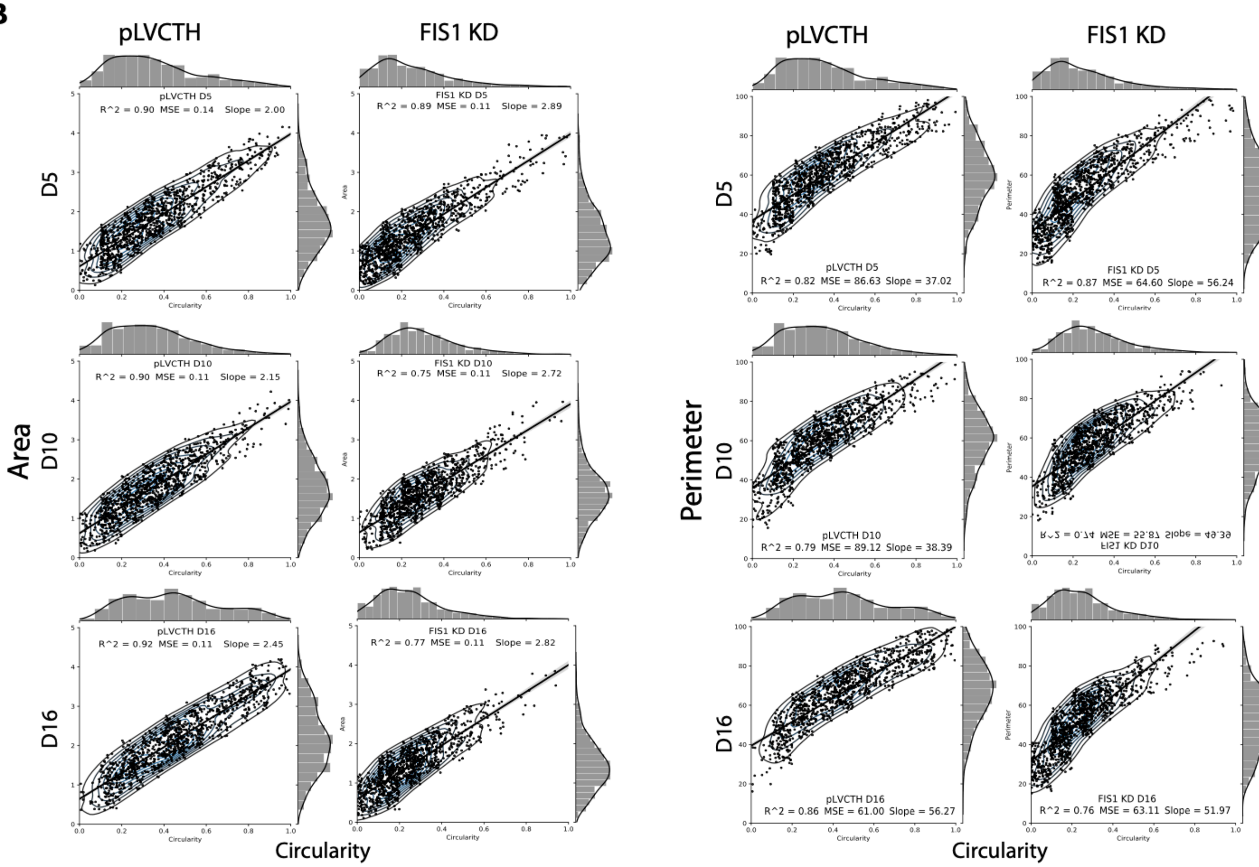
