## Supplemental Table I for "Exacerbation of mitochondrial fission in human CD34^+^ cells halts erythropoiesis and hemoglobin biosynthesis"

**Supplemental Table I: Primers used for qPCR**

| **Target gene** | **Organism** | **Accession number** | **Forward** | **Reverse** |
| --- | --- | --- | --- | --- |
| 18S rRNA | Homo sapiens | NR_003286.2 | Cgc tac act gac tgg ctc ag | Aaa ggg cag gga ctt aat caa c |
| Fis1 | Homo sapiens | NM_016068.2 | Agg cct taa agt acg tcc gc | Tgc cca cga gtc cat ctt tc |
| Mff | Homo sapiens | NM_001277061 | Ctt tcg cgc ttc tgc cct | Gag cca ctt ttg tcc ccc tg |
| Drp1 | Homo sapiens | NM_012063.2 | Gct cca gga cgt ctt caa ca | Ctt tcc gct gct ctg cgt tc |
| MiEF1 | Homo sapiens | NM_019008 | Cgg cct gcg gtg tga c | Gcc cga tag aga ctc agc ac |
| MiEF2 | Homo sapiens | NM_139162.3 | Ttg gag tta ggg cct gct tg | caa tgg tct gcc tgc gtc |
| Mfn1 | Homo sapiens | NM_033540.2 | Agt tgg agc gga gac tta gc | Atc gcc ttc tta gcc agc ac |
| Mfn2 | Homo sapiens | NM_014874.3 | Aat ctg agg cga ctg gtg ac | Ctc cac cag tcc tga ctt cac |
| Opa1 | Homo sapiens | NM_130831.2 | Cgg gaa ctt gac cgg aat ga | Cgc agc tgg aag gta gat gt |
| 18S rRNA | Mus musculus | NR_003278.3 | Ctg ccc tat caa ctt tcg atg gta g | Ccg ttt ctc agg ctc cct ctc |
| Fis1 | Mus musculus | NM_025562.3 | Tgt ggc caa gta gag acc tt | Tca aaa ttc ttc aga tcc tcc aca |
| Mff | Mus musculus | NM_029409.2 | Ggc act gaa aac acc acc ac | Cca act gct cgg ctc tct tc |
| Drp1 | Mus musculus | NM_152816.3 | Cgg ttc cct aaa ctt cac ga | Gca cca ttt cat ttg tca cg |
| Mfn1 | Mus musculus | NM_024200.4 | Att gcc aca agc tgt gtt cg | Cta ggg acc tga aag atg ggc |
